# Unlocking the Power of CXCR2 Inhibition to Overcome Gemcitabine Resistance in Pancreatic Cancer

**DOI:** 10.1101/2025.06.25.661585

**Authors:** Caitlin Molczyk, Reegan Sturgeon, Sugandha Saxena, Esther Johnsson, Rakesh Bhatia, Namita Bhyrabhatla, Sushil Kumar, Surinder K. Batra, Rakesh K. Singh

**Affiliations:** Department of Pathology and Microbiology, 985950, Nebraska Medical Center, Omaha, NE 68198-5900; Department of Biochemistry and Molecular Biology, University of Nebraska Medical Center, Omaha, NE, 68198-5845

## Abstract

Pancreatic ductal adenocarcinoma (PDAC) is the fourth leading cause of cancer-related mortality, characterized by intrinsic resistance to conventional therapies and limited effective treatment options. In this study, we investigated the role of the CXCR2 axis in PDAC therapy resistance. CXCR2, a chemokine receptor, is actively involved in inflammation, tumor angiogenesis, and metastasis. Our working hypothesis is that CXCR2 contributes to PDAC chemotherapy resistance. To test this, we generated gemcitabine-resistant (GemR) lines using T3M4 and CD18/HPAF (CD18) cell lines. Baseline expression of CXCL1, CXCL5, and CXCL8 ligands were higher in GemR cells compared to parental cells. Upon gemcitabine treatment, parental cells exhibited a greater increase in CXCL1 and CXCL8 expression than GemR cells. Further analysis in T3M4 cells revealed a dose- and time-dependent increase in CXCL1 and CXCL8 expression following gemcitabine exposure. Next, we assessed whether targeting CXCR2 could enhance the therapeutic response. We treated parental and GemR cell lines with gemcitabine in combination with the CXCR2 antagonist SCH 479833 (SCH 47). Notably, lower concentrations of gemcitabine combined with SCH 47 were more effective than higher concentrations of gemcitabine alone in GemR cell lines. In both parental and GemR xenograft models, combination therapy with Navarixin (a CXCR2 antagonist) and gemcitabine demonstrated superior antitumor and antimetastatic activity compared to either treatment alone. In conclusion, these findings highlight the critical role of the CXCR2 axis in PDAC therapy resistance. Targeting CXCR2 enhances gemcitabine efficacy, offering a potential therapeutic strategy to overcome resistance in PDAC.

## Introduction

Pancreatic ductal adenocarcinoma (PDAC) is the fourth leading cause of cancer-related deaths in the United States, with the incidence of new cases in 2025 almost equaling the estimated number of deaths (1). The factors affecting this grim forecast include a lack of surveillance and detection methods, nonspecific and vague symptomology of PDAC, and a lack of treatment options (2). In addition to being detected in the late stages of the disease, PDAC has a high degree of resistance to the chemotherapies available (3). This resistance is likely a significant factor in the mortality of the disease. There are many known ways cancer cells build resistance to chemotherapy, including ABC transporters, changes in metabolism (4), exosome production (5) and signaling pathways internally (6) and in the tumor microenvironment (TME) to interact with immune cells (7), endothelial cells (8), fibroblasts (9), and even other tumor cells (10). Though these tumor cells utilize many routes to overcome chemotherapy, the underlying mechanism(s) of resistance remain unclear(11).

Recent findings in cancer have implicated the role of chemokines in therapy resistance and metastasis (12–14). Chemokines have been classified into four main subfamilies: CXC, CC, CX3C, and XC. In normal tissues, chemokines are involved in the inflammatory responses (13, 15). CXC chemokines are involved in the recruitment of immune cells, specifically neutrophils, to the site of inflammation and subsequent regeneration of the damaged tissue (16, 17). However, during tumorigenesis, cancer cells are known to upregulate chemokines, but surprisingly, rather than suppressing the oncogenic activity of cancer cells, they help tumor progression instead (18, 19). We recently reported an increased expression of CXCR2 ligands in the supernatants of chemotherapy-treated breast cancer cells (20). Our observations were supported by clinical studies that showed elevated serum levels of CXCL6 and CXCL8 in patients with metastatic cancers (21, 22). Similarly, levels of CXCL1 and CXCL8 have been reported to increase after chemotherapeutic treatment (14, 23, 24). We furthermore observed that attenuation of the CXCR2 signaling axis sensitized mammary tumor cells towards chemotherapy both *in vitro* and *in vivo* (20). CXCR2 targeting in PDAC mouse models reduced metastasis, angiogenesis, and tumor volumes (25, 26). Litman-Zawadzka et al. reported that serum CXCL8 is a better biomarker for PDAC diagnosis and progression compared to CA19-9 (27). Together, these observations suggest that CXCR2 and its ligands are associated with tumor progression, therapy resistance, and metastasis. However, the precise role of the CXCR2 axis in therapy resistance has yet to be examined in PDAC.

Based on accumulating evidence and our previous findings, we hypothesized that the CXCR2 axis aids in PDAC chemotherapy resistance and targeting CXCR2 signaling will enhance therapeutic response. We generated Gemcitabine-resistant (GemR) variants of T3M4 and CD18/HPAF (CD18) cells and used it for our *in vitro* and *in vivo* analysis. Our data demonstrates the critical role of the CXCR2 axis in PDAC therapy resistance and its targeting sensitizes PDAC cells to gemcitabine, offering a potential therapeutic strategy to overcome resistance in PDAC.

## Methods and Materials

### Cell lines and reagents

PDAC human cell lines T3M4 and CD18 were maintained in Dulbecco’s Modified Eagle Medium (DMEM)(HyClone®, Thermo Scientific, UT). We supplemented the medium with 5% Fetal Bovine Serum (FBS) (Atlanta Biologicals, GA), L-Glutamine (MediaTech, VA), Vitamin solution (MediaTech), and Gentamycin (Gibco, Life Technologies, NY).

T3M4 and CD18 cell lines were cultured for three to seven days in increasing concentrations of gemcitabine. These cells were treated in this manner over six months (**Fig. 1A**). They then underwent a proliferation assay, the (3-(4,5-dimethylthiazol-2-yl))-2,5-Diphenyltetrazolium Bromide (MTT) assay, with the parent cell line as a control. With the analysis from Prism 7, we found the relative IC_50_. When comparing the IC_50_ of the parent and GemR variants, a ten-fold or higher increase compared to the parent cell line relative IC_50_ was considered resistant. MTT relative IC_50_ results are in **Fig. 1A**.

**Figure 1.**
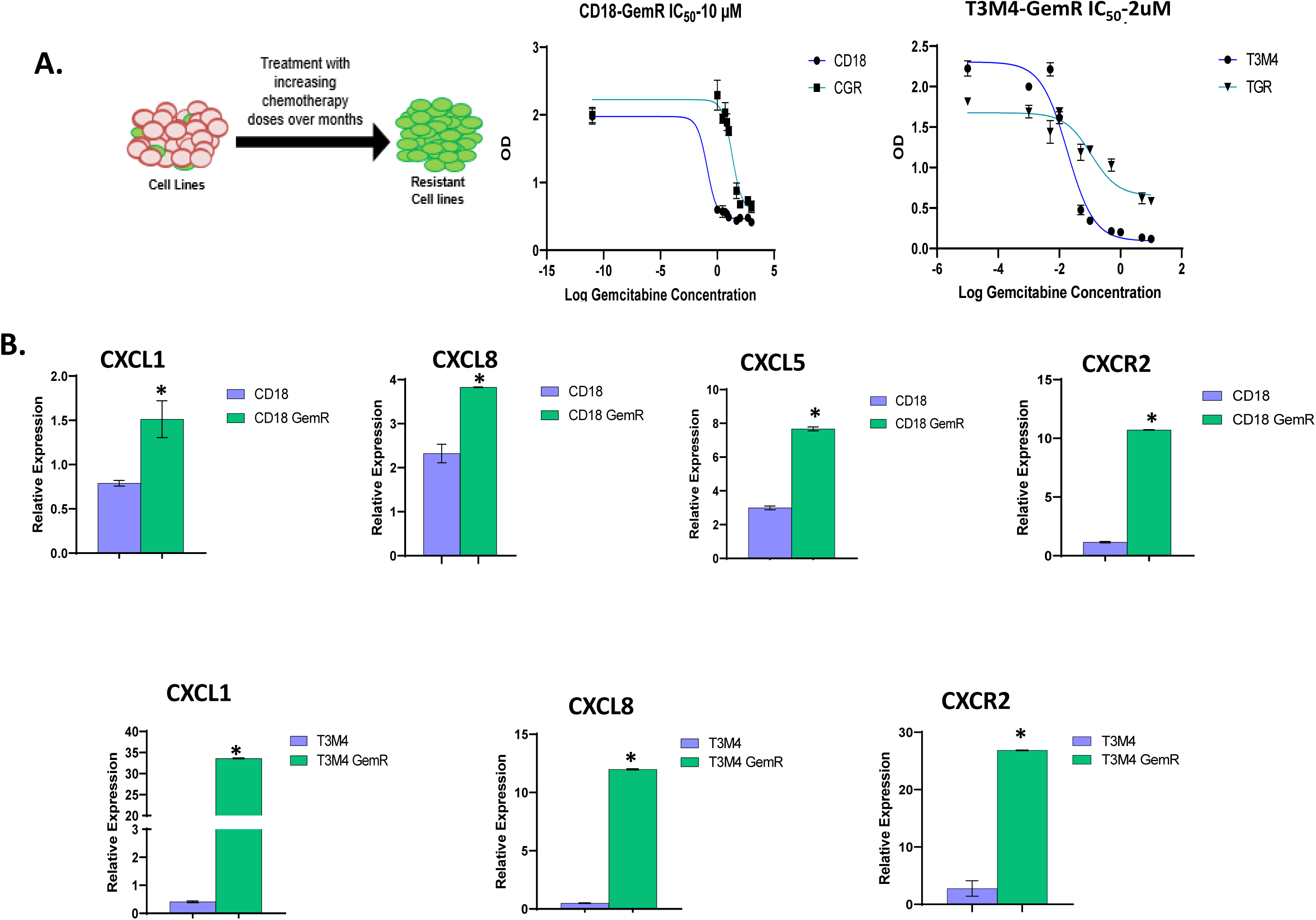
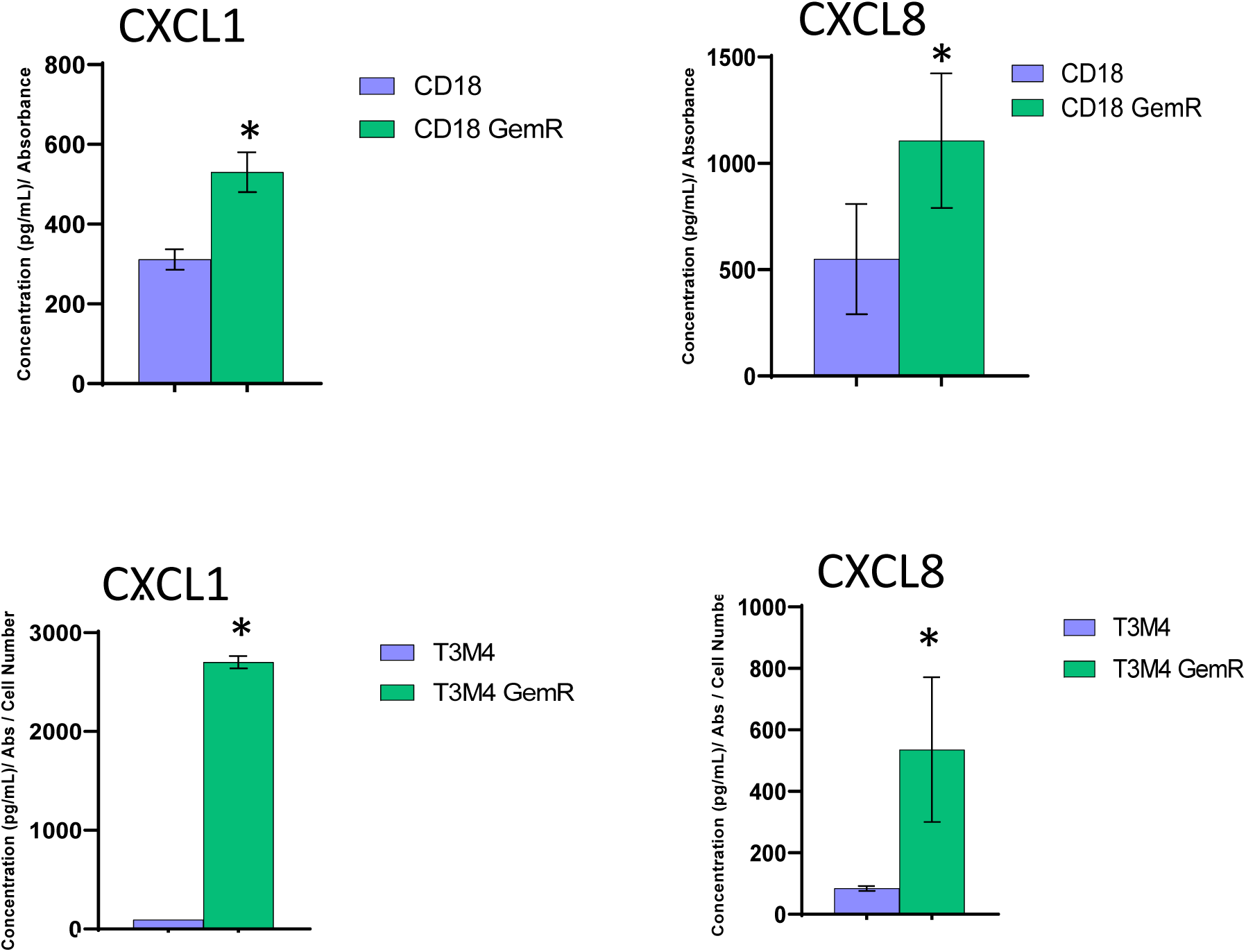
Gemcitabine-resistant cells express higher levels of CXCR2 and its ligands: **A.** The generation of GemR cell lines consisted of treating the parent cell line with increasing concentrations of Gem over several months until the cells reached a point where they could no longer grow in the higher concentration of Gem. IC for CD18-GemR cells (10uM) and T3M4-GemR cells (2uM) were ten-fold higher than the parental cells **B.** The mRNA expression of CXCR2 and its ligands was analyzed in CD18, T3M4, CD18-GemR, and T3M4-GemR cells using qRT-PCR. Gemcitabine-resistant cells express significantly higher levels of CXCR2 and its ligands CXCL1, 5, and 8. **C.** The secretion of CXCL1 and CXCL8 was analyzed in parents and gemcitabine-resistant tumor cells using ELISA, and a higher basal level of secretion was observed in gemcitabine-resistant tumor cells compared to parental cells. *p<0.01 is significantly different as compared to parental cells.

Small molecule CXCR2 antagonist SCH-52123 (Navarixin) was obtained from Schering-Plough Research Institute. Gemcitabine (Sagent and Meathial, NDC: 25021-234-10 and 71288-113-10) was obtained through the outpatient pharmacy at the Nebraska Medicine.

### mRNA expression Analysis

Total RNA was isolated using Trizol (Sigma) following the manufacturer’s protocol. Reverse transcription was performed with 1 µg RNA using Applied Sciences Reverse Transcription Supermix for RT-PCR. Quantitative real-time PCR (qRT-PCR) reactions were performed using iTaq Universal SYBR Green Supermix (Bio-Rad), using the QuantStudio 3 System. Primers used for the study are listed in **Table 1**. To analyze the qRT-PCR values, a housekeeping gene control (ribosomal protein large 13A (RPL13)) was used to normalize the C_t_ value for the relative expression of the gene of interest by [delta C_T_ = RPL13 C_t_ – Target gene C_t_]. The normalized C_t_ value was then taken (2^-delta^ ^C^_T_) * 10000 to obtain the relative expression. A melting curve analysis was performed to check the specificity of the amplified product.

**Table 1.**
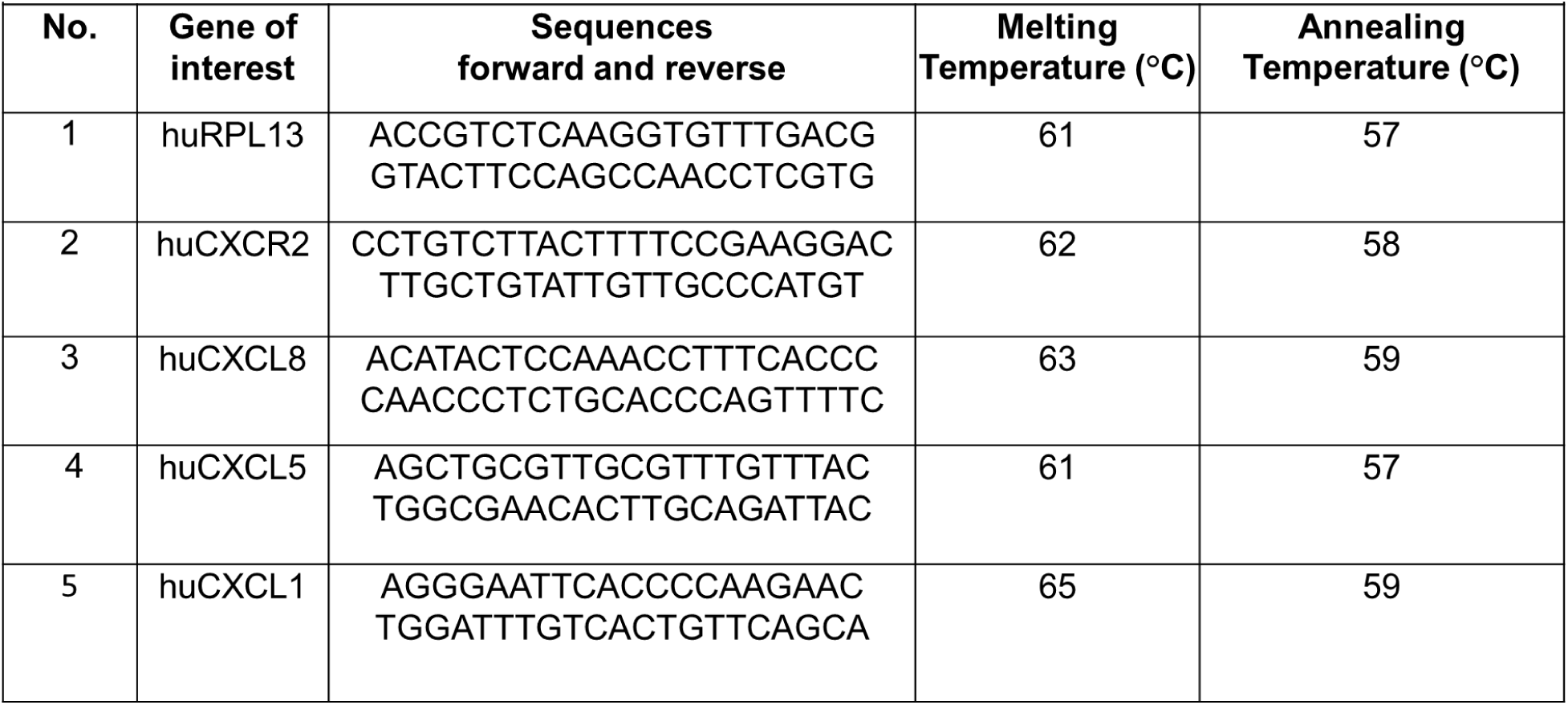
Primers used in this study.

### Enzyme-linked immunosorbent assay

Cells were seeded at densities ranging from 4 × 10^3^ to 2 × 10^6^ in a 96-well plate, 6-well plate, or 100 mm X 15 mm petri dish. Cells were treated with gemcitabine and/or Navarixin at various concentrations as depicted in the figures. The supernatants of these cells were collected for the enzyme-linked immunosorbent assay (ELISA). ELISAs for human (hu) CXCL1, huCXCL5, huCXCL8, muCxcl1, muCxcl5, muCxcl7 were performed using a Duoset kit (R&D Systems, Minneapolis, MN) according to the manufacturer’s protocol, using a Bio-Tek plate reader.

### Cell proliferation assay

Cells were plated at densities ranging from 3.5 to 5 × 10^3^ in 96-well plates. Plated cells were treated in triplicate with increasing concentrations of Gem (0, 0.001, 0.005, 0.01, 0.05, 0.1, 0.5, 1, 5, 10, 25, 50, 100, 500, 1000 µM) or CXCR2 antagonist (0, 5, 10, 25, 50, 100 µg/mL). Cells were treated for 72 hours with chemotherapy. After treatment with chemotherapy, MTT was added to each well. After an incubation of 4 hours, the supernatant was removed, and 100 µL of dimethyl sulfoxide (DMSO) was added to each well to lyse the cells. The absorbance of each well was read at 570_nm_ wavelength. The absorbances were used to analyze the percent inhibition of cell growth: [100 – (A/B) × 100], where A is the absorbance of the treated group and B is the absorbance of the control group, respectively. Anything over a 10% error was excluded from the statistical evaluation in the triplicate values.

### Gemcitabine/Navarixin combination treatment

For the combination treatment, plated cells were treated in triplicate with IC_50_ or half-IC50 concentrations of gemcitabine or Navarixin alone or in combination. The treatment incubation period was 72 hours, and MTT assay, mRNA analysis, and ELISA were performed as described earlier.

T3M4 cells were treated with 100 nM gemcitabine for different durations to examine duration-dependent effects. The samples were collected at 8, 12, 24, 48, and 72 hours, and MTT assay, mRNA analysis, and ELISA were performed as described earlier.

To determine the gemcitabine-induced concentration dependent changes in the expression of CXCR2 and its ligands, cells were treated with different concentrations of gemcitabine (0, 10, 50, 100, and 500 nM) for 72 hours and MTT assay and mRNA analysis were performed.

### Tumor growth and metastasis

Immunodeficient BALB/c or nude mice (6-8 weeks old) were purchased from the National Cancer Institute (Bethesda, MD). In accordance with approval from the American Association of Laboratory Animal Care and current regulations and standards of the US Department of Agriculture, Department of Health and Human Services, and National Institute of Health, the mice used for this study were maintained in facilities under specific pathogen-free conditions. All procedures performed in mice were in accordance with institutional guidelines and approved by the University of Nebraska Medical Center Institutional Animal Care and Use Committee (IACUC). CD18 and CD18-GemR cells (5 × 10^5^ cells in 30 µL HBSS) were orthotopically implanted. Seven days of post-inoculation, the mice were randomized into four treatment groups: control, Navarixin (5mg/Kg), Gemcitabine (25 mg/kg) alone, and a combination of Navarixin and Gemcitabine. Animals were treated daily with intraperitoneal injections. Primary tumors and spleen recovered from mice were fixed in formalin, embedded in paraffin, and processed for histopathological evaluation and immunohistochemistry. The livers were isolated and fixed in Bouin’s fixative for monitoring liver metastases macro- and microscopically.

### Hematoxylin and Eosin Staining (H & E)

To demonstrate the general histological architecture of tissue, we did regressive H & E staining. Hematoxylin demonstrates nuclear details, and the Eosin provides cytoplasmic and connective tissue details. The slides were stained in our Tissue Science Facility with an automated stainer Tissue-Tek® PrismaTM (Sakura, Torrance, CA). Briefly, 4-5 μm thick, formalin-fixed, paraffin-embedded tissue sections were stained with Hematoxylin (CV select HemaMax, BBC Biochemical, McKinney, TX) for 5 minutes and with Eosin (CV select Eosin Y, BBC Biochemical, McKinney, TX) for 90 seconds, then dehydrated from water, alcohol, and xylene with 30 second incubations steps. The slides were examined under Nikon Eclipse E800 light microscope after mounting with Permount.

### Immunohistochemistry/Immunofluorescence Analysis

Immunohistochemistry (IHC) or immunofluorescent (IF) staining was performed using 4-5 μm thick, formalin-fixed, paraffin-embedded tissue sections. After deparaffinization, antigen retrieval, and inhibition of non-specific binding, sections were probed with respective primary antibodies (Table 2) overnight at 4 °C. The following day, slides were washed and incubated with appropriate secondary antibodies. The ABC Elite Kit and DAB substrate kit (3, 3 Diaminobenzidine) (Vector Laboratories, Burlingame, CA) were used to detect Immunoreactivity per the manufacturer’s protocols. A positive IHC staining gave a brown color (cytoplasmic, nuclear, membrane, or all, depending on the location of the protein), while the nuclei were counterstained blue with Meyer’s Hematoxylin (Thermo Scientific, Fremont, CA). At least two independent observers evaluated IHC and IF staining. For quantitative evaluation, in each tissue section of the slides, positive cells were counted in five areas with significant staining at 200X magnification. Similarly, the number of positive endothelial cells was counted for blood vessels and lymphatics to evaluate microvessel density or lymphatic vessel density. Single or clusters of positive endothelial cells were considered a countable vessel, but the presence of blood cells or fibrin without any detectable endothelial cells was disregarded. All the above analyses were done, and the representative photomicrographs were captured with a Nikon Eclipse E800 microscope and its NIS-Elements BR 5.11.00 software (Nikon, Melville, NY).

**Table 2.** Antibodies used in this study.

| **Table 2. Antibodies used in this study** |  |  |  |  |  |
| --- | --- | --- | --- | --- | --- |
| **No.** | **Primary Antibody** | **Marker** | **Source** | **Catalog No.** | **Dilution** |
| 1 | Anti-Ki-67 | Cell Proliferation | Santa Cruz, TX, USA | sc-23900 | 1:100 |
| 2 | Anti-MPO<br>(Myeloperoxidase) | Neutrophils | Abcam, MA, USA | ab9535 | 1:100 |
| 3 | Anti-CC3<br>(Cleaved Caspase 3) | Apoptosis | Cell Signaling, MA, USA | asp175-5A1E-9664 | 1:200 |
| 4 | Anti- CD31<br>(Cluster of Differentiation 31) | Blood Vessels | Abcam, MA, USA | ab28364 | 1:200 |

### Masson Trichrome Staining

We used Masson Trichrome (MT) to stain the collagen in the tissue and differentiate it from muscle fibers and cytoplasm, which all normally get pink staining with routine H&E. 4-5 μm thick, formalin-fixed, paraffin-embedded tissue sections were stained with Masson Trichrome Stain Kit-Aniline Blue (BBC Biochemical, McKinney, TX) as per the manufacture’s protocol. In brief, staining with Aniline Blue for 2 minutes gives a blue color to the collagen, and using the other reagents in the kit the muscle fibers stain red and the cytoplasm stains pink. The slides were mounted with Permount and examined under the Nikon Eclipse E800 microscope. The amount of fibrosis (blue-stained collagen) was analyzed qualitatively and semi-quantitatively by the pathologist.

### Statistical analysis

The statistical analysis for all experiments was performed using the Prism 7 (GraphPad) software. The sample number and statistical method are indicated in the figure legends. Statistical significance was defined at p < 0.05. The error bars in the figures represent the standard error of the mean. When appropriate, the analyses included the two-tailed Student’s t-test, ANOVA, and Posthoc comparisons using Mann-Whitney tests with a Bonferroni adjustment.

## Results

### GemR cells express higher levels of CXCR2 ligands

The earlier report from our laboratory showed baseline differences between parent and chemotherapy-resistant cell lines in the CXCR2 axis gene expression in breast cancer (20). We examined whether therapy-resistant PDAC utilizes a similar pathway. We used CD18, T3M4, and their gemcitabine-resistance variants and analyzed the expression of CXCR2 and its ligands. T3M4-GemR and CD18-GemR cell lines have increased CXCL1, 5, and 8 mRNA and CXCR2 expression compared to the parental cell line (**Fig. 1B**). While all expression data was statistically significant, CXCR2 mRNA expression was significantly higher in GemR cell lines with each having an almost 10-fold increase when compared to the parental cell lines (**Fig. 1B**). We observed similar results and protein levels using ELISA (**Fig. 1C**). This data suggests upregulated expression of basal CXCR2 and its ligands following development of gemcitabine resistance.

### Gemcitabine differentially regulates expression of CXCR2 and its ligands’

Next, we elucidated whether gemcitabine treatment could induce CXCR2 ligand expression. The parental and GemR variants were treated with gemcitabine, and CXCR2 mRNA expression and protein secretion were analyzed. In the parental CD18 and T3M4 cells, gemcitabine treatment significantly enhanced CXCR2 and its ligands CXCL1 and 8 mRNA expression (**Fig. 2A**). We did not observe similar induction in CD18-GemR and T3M-GemR cells (**Fig. 2A**). In contrast, we observed the downregulation of CXCL1 and CXCL8 in T3M4-GemR cells as compared to the T3M4 parent cells (**Fig. 2A**). We evaluated protein secretion for these genes by ELISA. The results suggested increasing secretion of CXCL1 and CXCL8 protein in gemcitabine-treated parent cell lines (**Fig. 2C**); however, we did not observe any change in GemR cells treated with gemcitabine. This data suggests that gemcitabine treatment significantly upregulates CXCR2 and its ligands expression in PDAC parent cell lines.

**Figure 2.**
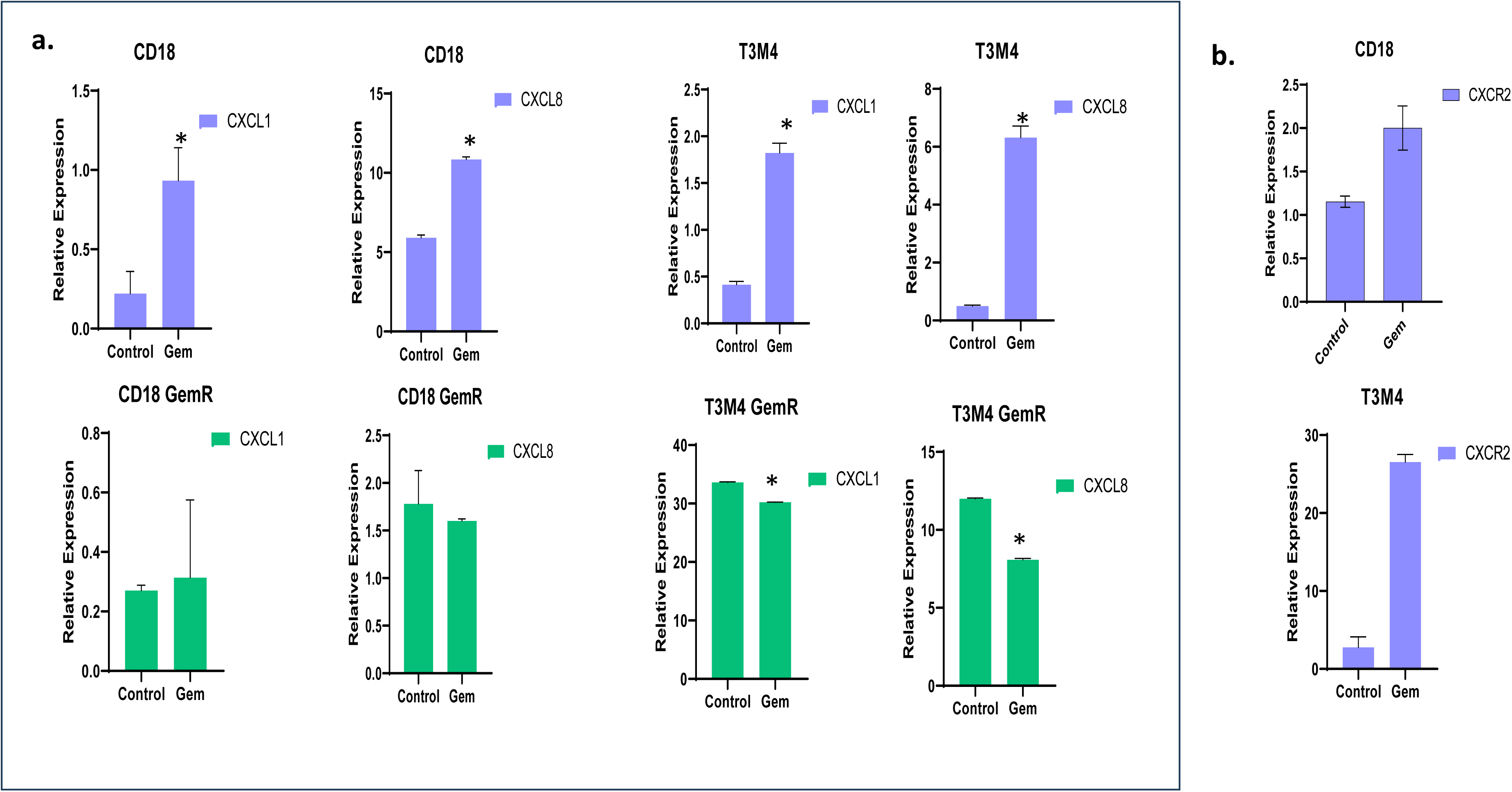
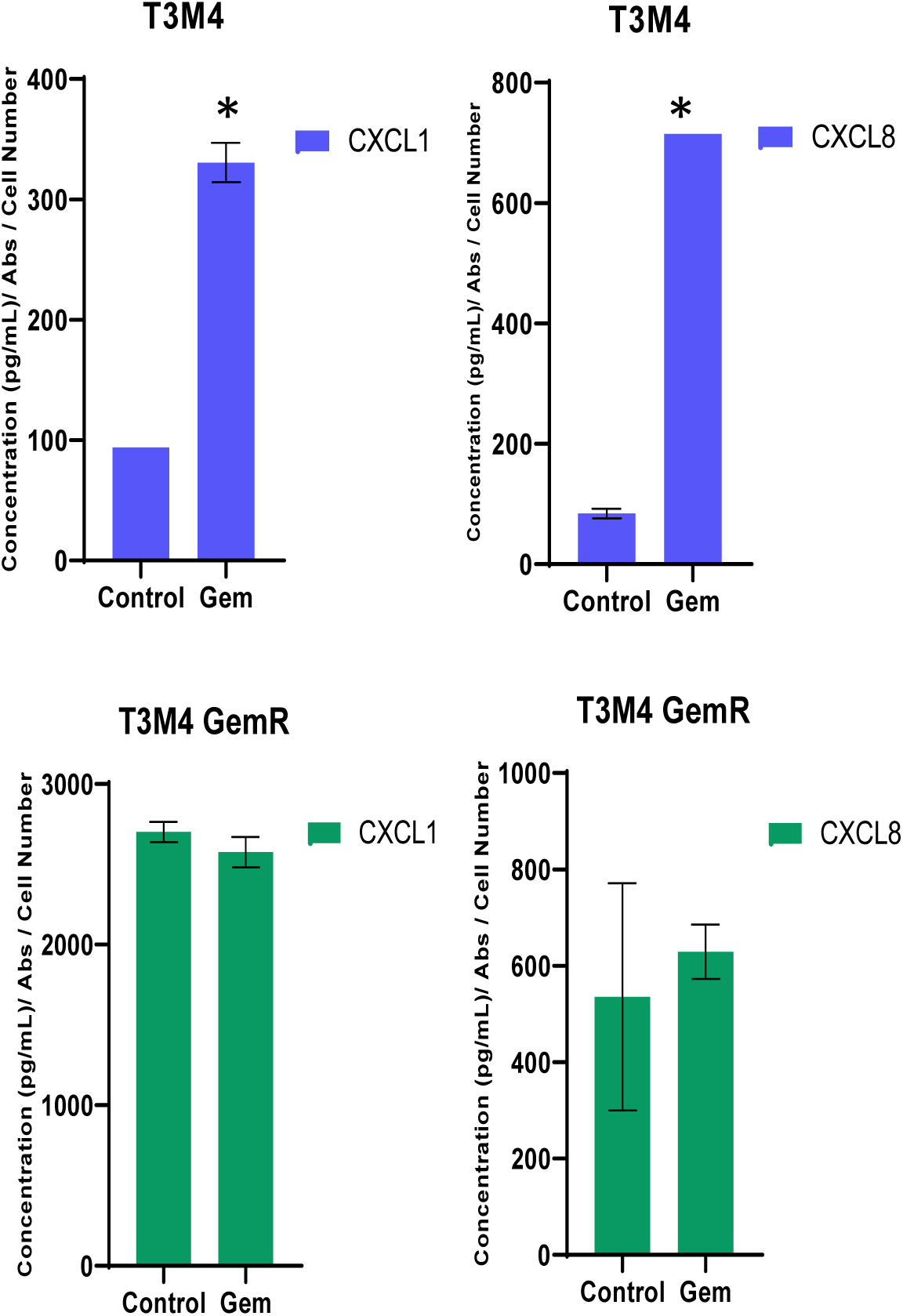
Gemcitabine differentially regulates the expression of CXCR2 and its ligands in a concentration and duration-dependent manner: **A. Expression of** CXCR2 and its ligands was analyzed in parental and chemotherapy-resistant cell lines following treatment with gemcitabine using qRT-PCR. Increased expression of CXCR2 ligands was observed in gemcitabine-treated parental cells, whereas no upregulation of CXCR2 ligands was observed following gemcitabine in GemR cells (**Aa**). The CXCR2 mRNA expression was also upregulated following gemcitabine treatment (**Ab**). **B.** The secretion of CXCL1 and CXCL8 was analyzed in parents and gemcitabine-treated tumor cells using ELISA. Gemcitabine treatment enhanced CXCL1 and CXCL8 secretion in parental cells. Resistant cells had a higher basal level of secretion; however, gemcitabine did not further enhance the CXCL1 and CXCL8 secretion. *p<0.01 is significantly different from untreated controls.

In the next set of experiments, we examined whether gemcitabine-induced CXCR2 ligand expression is concentration and/or duration-dependent. T3M4 cells were treated with different concentrations of gemcitabine for 72 hrs, and CXCR2 and associated ligands’ expressions were analyzed. We observed increased mRNA expression of CXCR2 and its ligands, CXCL1 and CXCL8, in a concentration-dependent manner (**Fig. 3A**). Similarly, we observed a concentration-dependent increase in the secretion of CXCL1 and CXCL8 in gemcitabine-treated T3M4 cells (**Fig. 3 A**). Next, CXCR2 and its ligands were analyzed to determine whether their modulation following gemcitabine treatment is dependent on the duration of the treatment. T3M4 cells were treated with gemcitabine (100 nM) for 8, 12, 24, 48, and 72 hours, and mRNA expression and protein secretion were analyzed. We observed duration-dependent modulation in the mRNA expression of CXCR2 and its ligands (**Fig. 3B**). Similarly, we observed a duration-dependent increase in CXCL1, CXCL5, and CXCL8 secretion in gemcitabine-treated T3M4 cells (**Fig. 3B**). Together, these data demonstrate that gemcitabine treatment upregulates the expression of CXCR2 and its ligands in PDAC cells.

**Figure 3.**
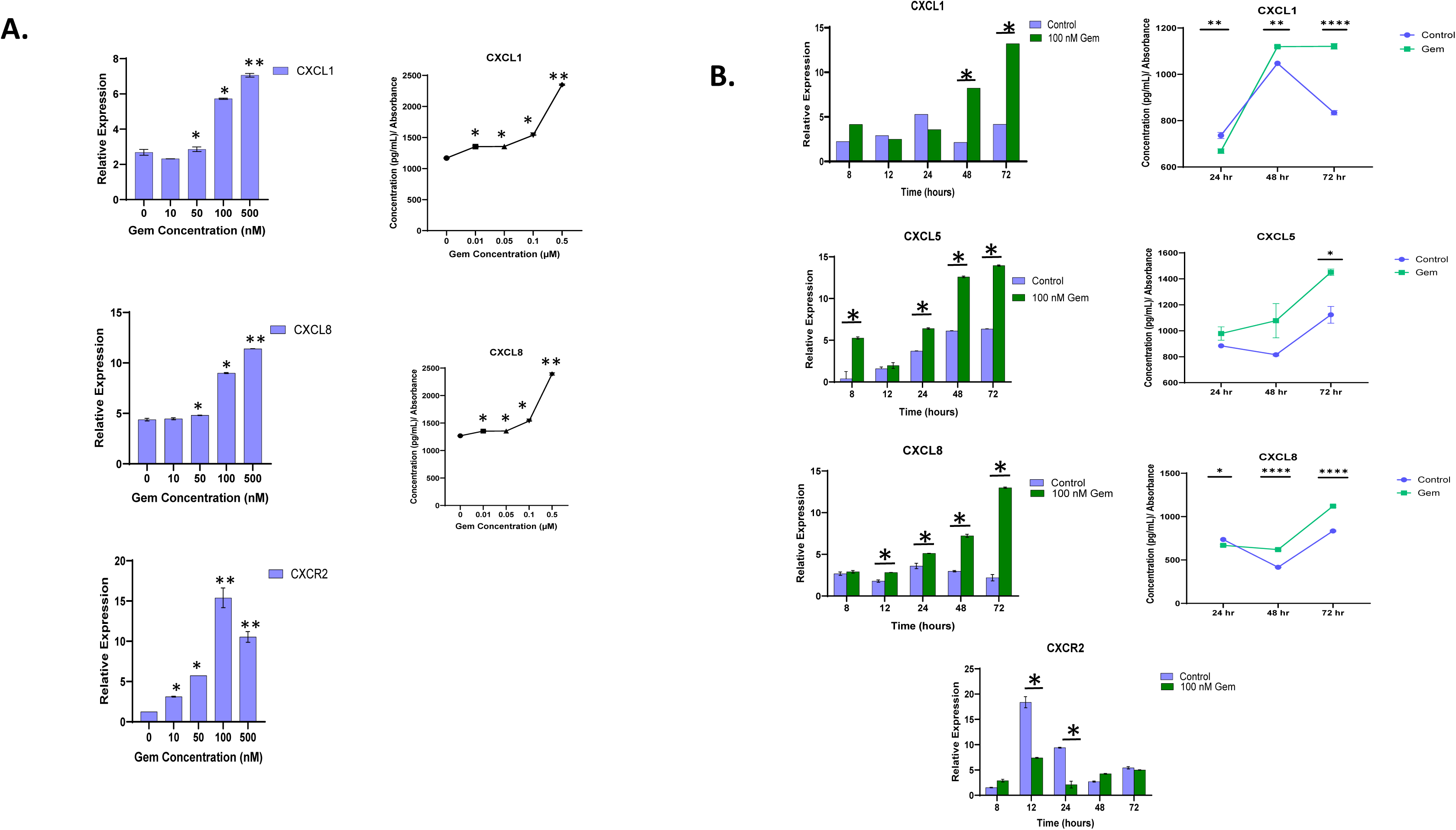
Concentration- and duration-dependent expression of the CXCR2 and its ligands. A. The T3M4 cell line was treated with increasing concentrations of Gem (0-500 nM) for 72 hours, and the CXCR2 and its ligand expression were assessed using qRT-PCR. There is a concentration-dependent increase in the CXCL1, CXCL8, and CXCR2 expression following gemcitabine treatment. **B.)** To analyze duration-dependent response, T3M4 cells were treated with 100 nM of gemcitabine for 8, 12, 24, 48, and 72 hours, and CXCR2 and its ligands expression were examined using qRT-PCR. The mRNA expression showed increases in the expression of CXCL1, CXCL5, CXCL8, and CXCR2 over the period. *p<0.05, **p<0.01, ****p<0.001 is significantly different from untreated controls.

### Inhibition of CXCR2 signaling enhanced gemcitabine-mediated antitumor response in GemR cells

We analyzed whether a CXCR2 antagonist can enhance the antitumor response of gemcitabine in therapy-resistant cells. CD18, T3M4, and their gemcitabine-resistant variants were treated with different concentrations of Navarixin alone for 72 hrs, and the proliferative response was monitored. We observed that Navarixin alone inhibited tumor cell proliferation in parent and GemR cells in a concentration-dependent manner (**Fig. 4A**). In the next set of experiments, we used sub-optimal concentrations of Navarixin, gemcitabine alone, or in combination. Our data suggest that the inhibition of CXCR2 signaling using small molecule antagonist, Navarixin, enhanced the therapeutic response of gemcitabine in parental and gemcitabine-resistant cells (**Fig. 4B**). The effectiveness of CXCR2 antagonism was more pronounced in gemcitabine resistant cell lines as compared to parental cells (**Fig. 4B**). These data demonstrate that inhibition of CXCR2 signaling potentiates an antitumor response in therapy-resistant tumor cells.

**Figure 4.**
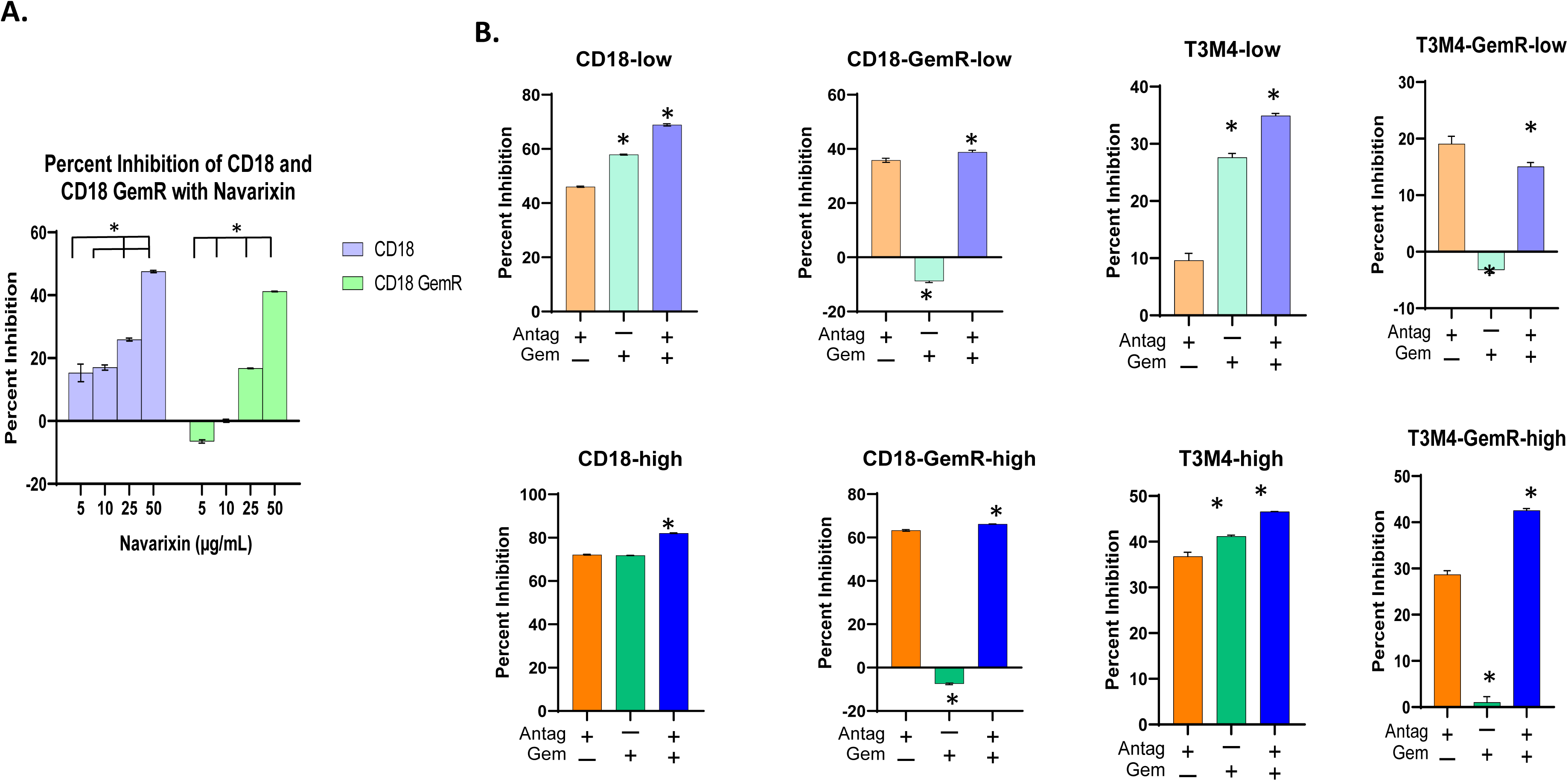
Inhibition of CXCR2 signaling enhanced gemcitabine-mediated antitumor response in gemcitabine-resistant cells. **A.** CD18 and CD18-GemR cells were treated with different concentrations of Navarixin for 72 hours, and inhibition in cell proliferation was assessed using the MTT assay. **B.** CD18, T3M4, and their GemR variants (CD18-emR and T3M4-GemR) were treated with varying combinations of low and higher doses of Navarixin and gemcitabine for 72 hours. CD18-low/CD18-GemR-low combination included 25 uM Navarixin plus 75nM gemcitabine, whereas CD18-high/CD18-GemR-high combination included (50 uM Navarixin plus 150nM gemcitabine. Similarly, for T3M4 cells low combination included 25 uM Navarixin and 25 nM gemcitabine, whereas the high combination included 50 uM Navarixin plus 50 nM gemcitabine. Inhibition in cell proliferation was analyzed using the MTT assay. *p<0.01 is significantly different from controls.

### CXCR2 antagonism enhanced gemcitabine antitumor and antimetastatic responses in therapy-resistant tumors

For these studies, we used a xenograft model of orthotopic implantation of CD18, and CD18-GemR cell lines. Seven days after the tumor implantation, animals were randomized to receive Navarixin, gemcitabine alone, or a combination twice weekly intraperitoneally. The treatment period for the mice injected with the GemR cell line was shortened by seven days due to the high tumor burden in the control group (**Fig. 5A**). The parent cell line-injected mice were treated for four weeks, whereas the GemR cell line-injected mice underwent treatment for three weeks. In CD18 parent cell implanted animals, there was a significant difference in the tumor size in animals treated with a combination of gemcitabine and Navarixin (**Fig. 5B**). We did not observe any difference in animals treated with Navarixin alone as compared to the control group. The gemcitabine-treated group had smaller tumors (non-significant) compared to the control group (**Fig. 5B**). However, results from CD18-GemR cell implanted animals demonstrate that Navarixin and gemcitabine alone significantly inhibited tumor growth, indicating the combination of Navarixin with gemcitabine was significantly effective (**Fig. 5B**). These data demonstrate that inhibition of CXCR2 signaling enhanced antitumor responses in gemcitabine-resistant tumors.

**Figure 5.**
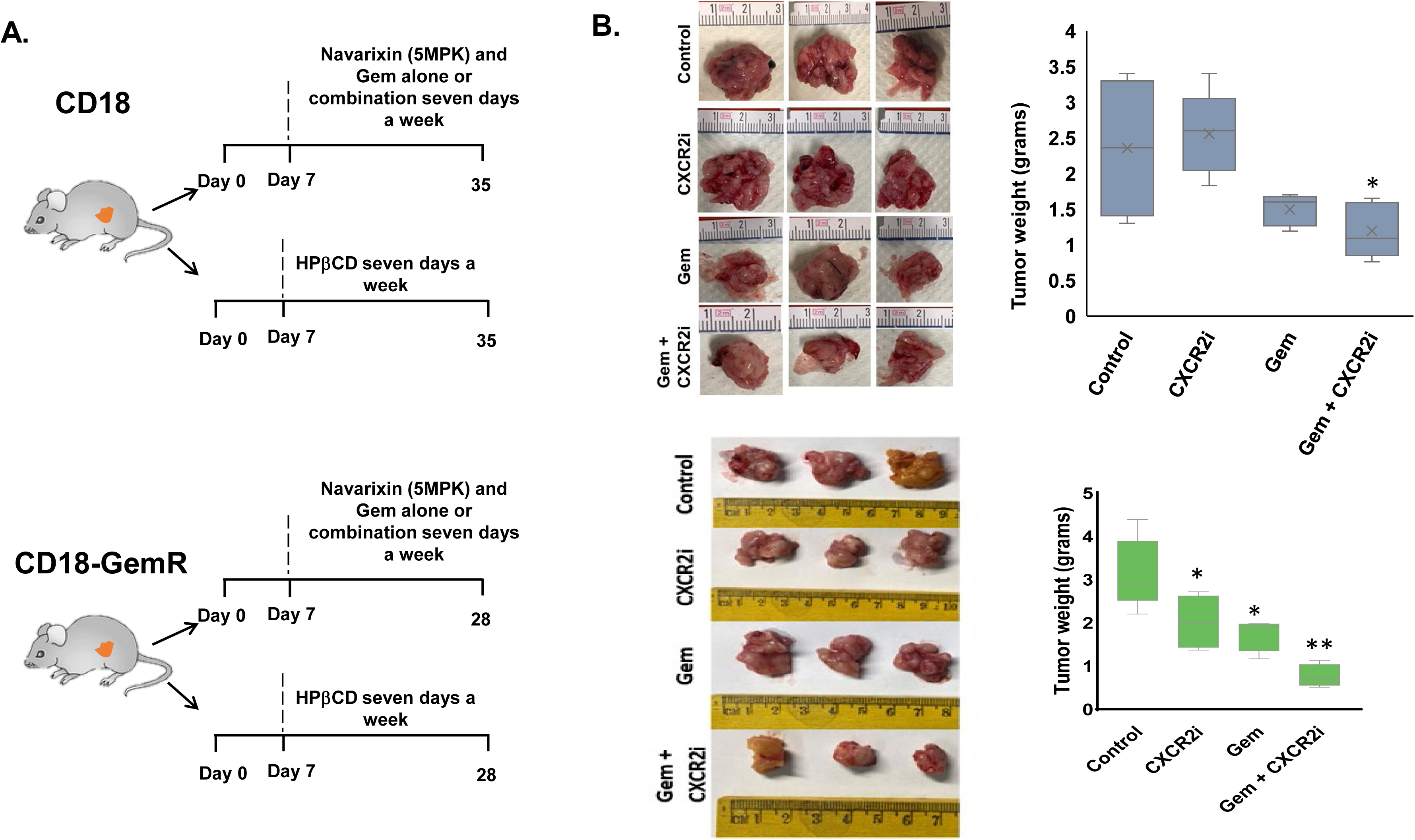
CXCR2 antagonism enhanced gemcitabine antitumor responses in therapy-resistant tumors. *In vivo* results from CD18 and CD18 GemR orthotopic injections (n=5 mice per cell line per treatment group). The animals were treated as described, and tumors were harvested as an end-point analysis. Tumor weight was compared between the control and treated groups. The values are average tumor weight ± SEM. *p<0.05, p**<0.01 is significantly different from controls.

Next, we analyzed the tumor sections for overall necrosis, infiltrating leukocytes, and micro-metastases using the H&E slides. Necrosis in the CD18-GemR tumors was highest in the combination treatment (**Fig. 6A**). Compared with the control, all treatment groups had higher immune infiltrates (**Fig. 6A**). We used Ki67 and CC3 immunostaining to evaluate *in situ* tumor cell proliferation and apoptosis, respectively. The cell proliferation was lowest in the combination-treated tumors, with the Navarixin-treated group being the second lowest (**Fig. 6A**). CC3^+^, apoptotic cells, did not change significantly between groups (**Fig. 6A**). Furthermore, the Masson trichrome staining evaluated the fibrosis in each tumor. We observed an increasing amount of fibrosis in the Navarixin- and combination-treatment groups compared with control and gemcitabine treatment (**Fig. 6A**).

**Figure 6.**
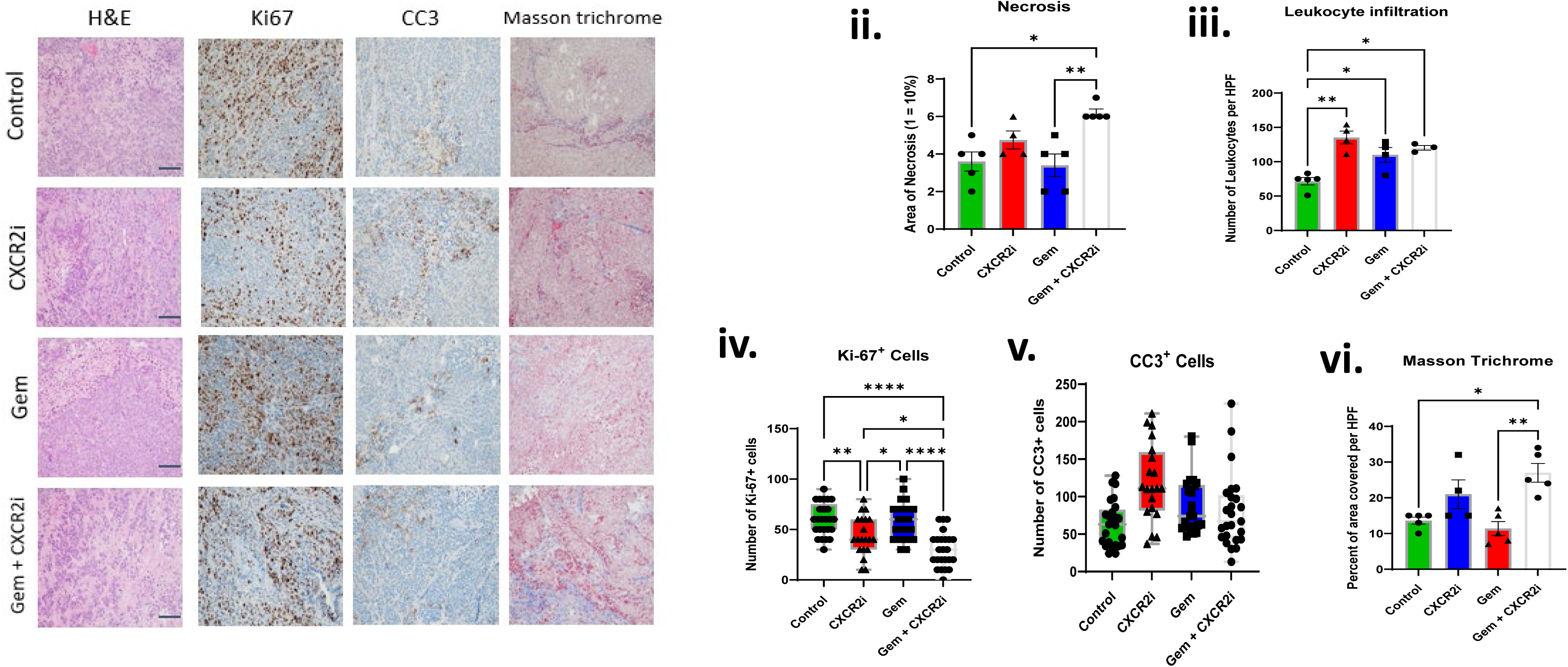
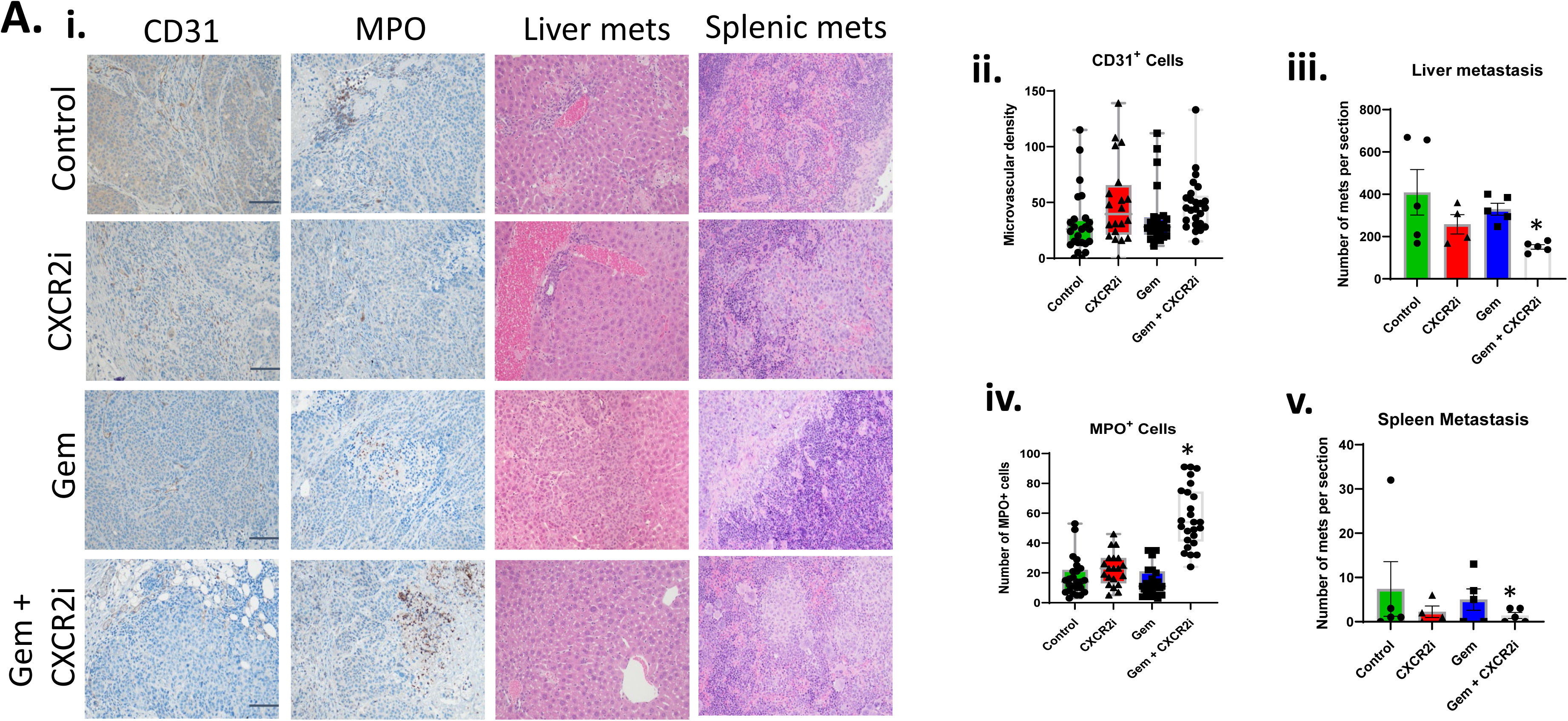
CXCR2 targeting modulated *in situ* proliferation, angiogenesis, leukocyte recruitment, and inhibited metastasis. The treatment and control tumors were analyzed for *in situ* cell proliferation (Ki-67), apoptosis (CC3), necrosis, leukocyte infiltration, and fibrosis (Masson Trichrome). The values are the average number of cells/areas ± SEM. C. Tumors’ angiogenesis (CD31), neutrophil infiltration (MPO), and liver and spleen metastasis were analyzed. *p<0.05 is significantly different from the control group. *p<0.05, **p<0.01, ****p<0.001 is significantly different from untreated controls.

In the next set of experiments, we analyzed neovascularization, neutrophil infiltration, and the extent of liver and spleen micro-metastasis in treated and control tumors. We found higher variability in all treatment groups, though there was no significant difference in the extent of neovascularization (**Fig 6Bii**) in various treatment groups. Compared to the control, there was no significant change in the number of MPO^+^ neutrophils in the gemcitabine and Navarixin alone treated tumors. However, the combination-treated group had a significant increase in neutrophil infiltration (**Fig. 6Biv**). These cells were in the larger necrotic/apoptotic regions of the tumors.

In the next set of experiments, we examined the efficacy of different treatment regimens on liver and spleen metastasis. There was a lower number of micrometastatic lesions in the liver and spleen in the Navarixin-treated group as compared to the control and gemcitabine alone-treated groups (**Fig. 6Biii + v**). More importantly, the combination treatment of Navarixin and gemcitabine significantly reduced the number of micro-metastases in the liver and spleen (**Fig. 6Biii + v**). Together, these data suggest that targeting CXCR2 enhances therapeutic responses in therapy-resistant tumors.

## Discussion

The urgency of understanding PDAC chemotherapy resistance is escalating as the five-year survival rate has had minimal improvement over the last ten years and remains around 10% (28) and alternative targetable pathways are warranted. In this study, we demonstrate that the CXCR2 axis is crucial for resistance in PDAC. In our *in vitro* modeling using T3M4 and CD18, we observed an upregulation of the CXCR2 axis in the derived resistant cell lines. Our findings and other reports suggest that chemotherapy resistance utilizes the CXCR2 axis (12, 29–33). We also found that PDAC parent cell lines have inducible CXCR2 expression via increasing chemotherapy dose and treatment duration. Sharma et al. found that in breast cancer cells, the chemotherapy-resistant cell lines had higher expression of the CXCR2 axis than the parent cell lines (20). In melanoma cell lines, Sharma et al. also found that the more aggressive melanoma cell lines had higher CXCR2 expression (34).

Using a small molecule CXCR2 antagonist in chemoresistance and chemosensitivity in PDAC is a novel finding. Earlier reports have demonstrated that the CXCR2 axis aids in many pro-tumor capacities, including angiogenesis (26, 35), metastasis (26, 33, 36), and chemotherapy resistance (8, 29, 33, 37) in different tumor types. Previous studies from our laboratory also suggest the ability of CXCR2 inhibition to re-sensitize tumors to chemotherapy with a CXCR2 antagonist or synergism between the CXCR2 antagonist and the chemotherapy (14). In CRC, Nav-treated mice had decreased tumor growth (38). Several groups found that the CXCR2 antagonism sensitized the tumors to chemotherapy (39, 40). Our data do not suggest re-sensitization of resistant cell lines to chemotherapy (40) or multidrug resistance (41); however, we report the ability of the antagonist alone to target the GemR cells therapeutically. This novel finding suggests that chemotherapy and the CXCR2 antagonist can target the sensitive tumor cells, while the antagonist can effectively target the chemotherapy-resistant cells.

Our present data demonstrated that targeting CXCR2 enhanced therapeutic responses in therapy-resistant tumors. Navarixin alone or in combination with gemcitabine significantly inhibited tumor growth and metastasis. We did not observe Navarixin alone having a significant effect on tumors derived from parental cells. Earlier reports from our laboratory and others have demonstrated the role of the CXCR2 axis in PDAC growth, angiogenesis, and metastasis (25, 26, 42, 43). The combination treatment with Navarixin and gemcitabine was most effective in both parental and gemcitabine-resistant tumors. Similarly, we observed a significant decrease in in situ cell proliferation, necrosis, and leukocyte infiltration in the combination treatment group. However, the observed enhanced antitumor and antimetastatic activity did not show an additive or synergistic response.

We observed decreased liver and spleen micrometastases in the Navarixin and gemcitabine combination treatment group. CXCR2 and its ligands have been shown to play an important role in metastasis (44, 45), and blocking this pathway has decreased metastases (13, 45, 46). Blocking the CXCR2 axis with an antagonist alone or in combination effectively decreases metastasis-promoting tumor cells or their ability to metastasize in our study. However, we did not observe a significant difference in the Navarixin alone treatment compared to the control group. We did not observe any significant difference in the extent of neovascularization. Interestingly, there was a significant increase in the number of infiltrating neutrophils in the Navarixin plus gemcitabine combination treatment compared to either treatment alone. It is likely due to the significant amount of necrosis with massive neutrophil infiltration in these tumors.

Unexpectedly, we observed significant antitumor activity of gemcitabine in gemcitabine-resistant tumors. Earlier reports have suggested that low-dose chronic gemcitabine treatment enhanced the immunogenicity of tumor cells and activated Natural Killer (NK) cells targeting tumor cells via chemoimmunotherapy (47, 48). The activation of NK cells is linked to chemotherapy, increasing the expression of NKG2D ligands in cells (47), which removes their invisibility cloak to the immune system. In our laboratory, we have seen that therapy-resistant breast cancer cells upregulate the expression of the murine NKG2D ligands Rae1 and Mult1 (14). We observed upregulation of human NKG2D ligands ULBP1, 2, 3, 4, 6, and MICB in the gemcitabine-resistant cells (**Supplemental Fig. 1**), which may have been susceptible to gemcitabine alone therapy due to increased NK cell activation.

In conclusion, the present study highlights the importance of CXCR2 signaling in PDAC therapy resistance and demonstrates that targeting this signaling may help reduce the problem of resistance in cancer. Chemoresistance is nuanced, especially in PDAC, due to the increased intrinsic and extrinsic resistance in this tumor and the tumor microenvironment. Our data demonstrates that CXCR2 targeting combined with gemcitabine enhanced antitumor and antimetastatic responses in therapy-resistant tumors, suggesting it to be an effective avenue for future PDAC therapeutic management.

## Acknowledgement

This work was partly supported by grants R01CA228524, Cancer Center Support Grant (P30CA036727) from the National Cancer Institute, and National Institutes of Health (R.K.S.), and Caitlin Molczyk is supported by a T32CA009476 Eppley Institute Cancer Biology Training Grant from the National Cancer Institute. We thank Samuel M. and Janel L. Cohen, Distinguished Professorship of Pathology and Microbiology Fund (R.K.S.), for their generous support.

---

**Supplemental Figure 1.**
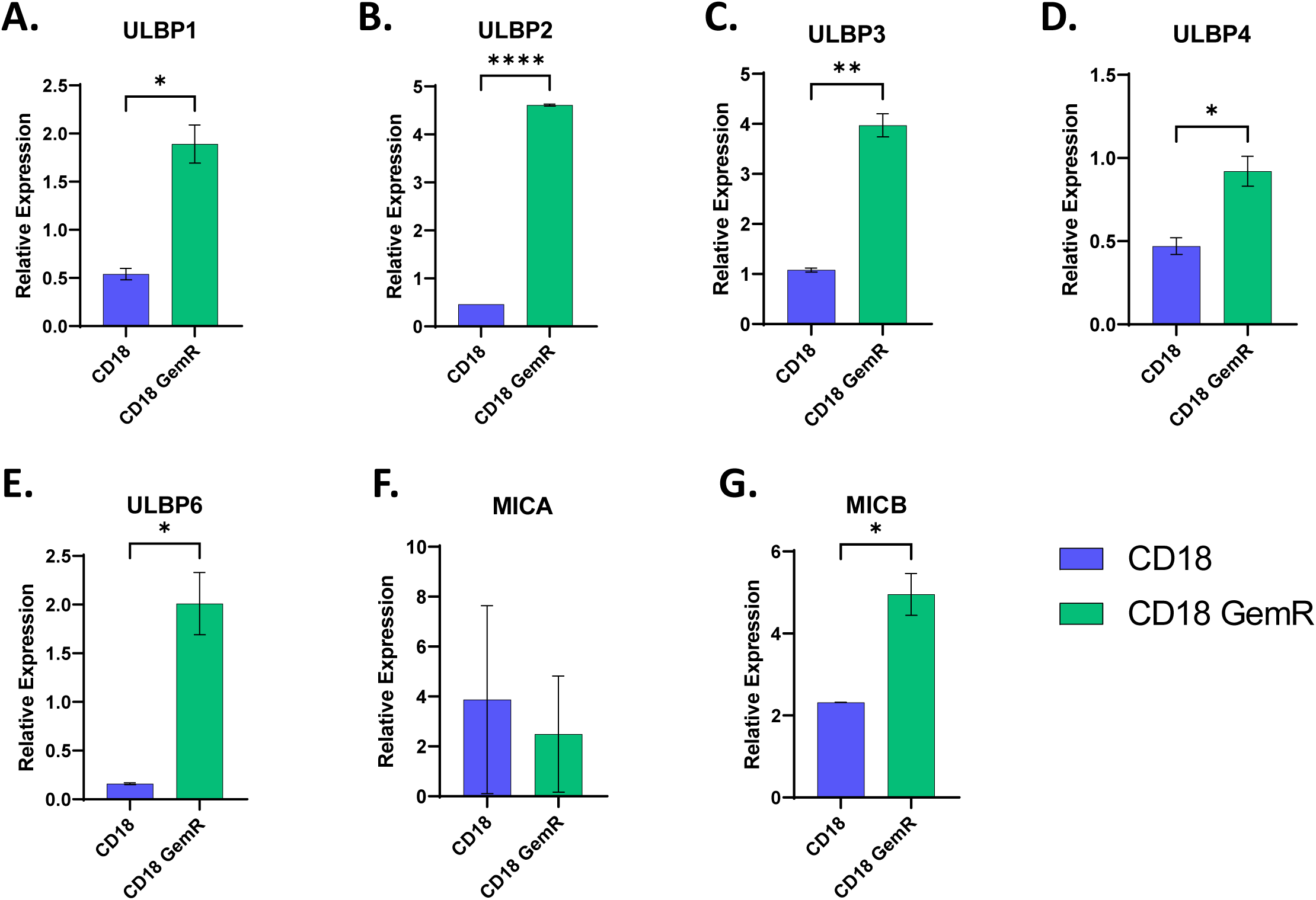
CD18 and CD18 GemR cells *in vitro* were analyzed for the expression of human NKG2D ligands, ULBP1-6, MICA, and MICB.

## Notes

**Conflict of Interest**: The authors declare no potential conflicts of interest.

### Competing Interest Statement

The authors have declared no competing interest.

